# Stress-induced changes in the S-palmitoylation and S-nitrosylation of synaptic proteins

**DOI:** 10.1101/408468

**Authors:** Monika Zareba-Koziol, Anna Bartkowiak-Kaczmarek, Izabela Figiel, Adam Krzystyniak, Tomasz Wojtowicz, Monika Bijata, Jakub Wlodarczyk

## Abstract

The precise regulation of synaptic integrity is critical for neuronal network connectivity and proper brain function. Essential aspects of the activity and localization of synaptic proteins are regulated by posttranslational modifications. S-palmitoylation is a reversible covalent modification of the cysteine with palmitate. It modulates affinity of the protein for cell membranes and membranous compartments. Intracellular palmitoylation dynamics are regulated by other posttranslational modifications, such as S-nitrosylation. Still unclear, however, are the ways in which this crosstalk is affected in brain pathology, such as stress-related disorders. Using a newly developed mass spectrometry-based approach (<u>Pa</u>lmitoylation And <u>N</u>itrosylation <u>I</u>nterplay <u>Moni</u>toring), we analyzed the endogenous S-palmitoylation and S-nitrosylation of postsynaptic density proteins at the level of specific single cysteines in a mouse model of chronic stress. Our results suggest that atypical mechanism of crosstalk between the S-palmitoylation and S-nitrosylation of synaptic proteins might be one of the major events associated with chronic stress disorders.

## Introduction

Synaptic plasticity is a dynamic process that is driven by molecular changes at individual synapses that lead to the rapid and recurrent rewiring of neuronal circuitry (Amtul and Atta-Ur-Rahman, 2015, Holtmaat and Svoboda, 2009). Each synapse contains thousands of different proteins, including receptors, signaling molecules, scaffolding proteins, and cytoskeleton components that are directly involved in synaptic transmission (Bayés et al., 2011, Xu, 2011). Understanding the mechanisms by which functions of synaptic proteins are regulated is crucial for elucidating the molecular basis of synaptic plasticity.

Synaptic proteins are regulated by a wide range of posttranslational modifications (PTMs), including *S*-palmitoylation (S-PALM) and *S*-nitrosylation (S-NO). These chemical modifications affect the properties of target molecules, resulting in the modulation of synaptic function and plasticity (Karve and Cheema, 2011, Okamoto et al., 2014, Yokoi et al., 2012). Recent evidence suggests that S-PALM may play a key role in regulating various synaptic proteins, including receptors, and ion channels (el-Husseini and Bredt, 2002, Fukata and Fukata, 2010, Fukata et al., 2013). *S*-palmitoylation refers to the addition of palmitoyl moieties to selected cysteine via thioester bonds, which increases protein hydrophobicity and the affinity for plasma membranes (Linder and Deschenes, 2007, Tortosa and Hoogenraad, 2018). One of the key features of S-PALM is its full reversibility, which makes it an exceptional mechanism for the rapid spatiotemporal control of intracellular and extracellular signaling. *S*-palmitoylation modulates multiple aspects of synaptic protein activity, including trafficking, conformational changes, stability, protein-protein interactions, and other PTMs (Fukata et al., 2013, Lussier et al., 2015, Conibear and Davis, 2010). The regulation of S-PALM is mediated by two types of enzymes: (*I*) protein acyltransferases (PATs) that attach the palmitate group to protein side chains and (*II*) acyl-protein thioesterases (APTs) that depalmitoylate proteins that hydrolyze thioester bonds (Greaves and Chamberlain, 2011, Salaun et al., 2010, Noritake et al., 2009, Won et al., 2018, De and Sadhukhan, 2018). In addition to enzymatic control, the dynamics of S-PALM are regulated by other protein modifications that occur in the target protein, such as phosphorylation and S-NO (Gauthier-Kemper et al., 2014, Salaun et al., 2010). Recently, the S-NO-dependent regulation of S-PALM dynamics was reported. The covalent modification of a protein’s cysteine thiol by a nitric oxide group appears to directly compete for cysteine residues with the palmitate or even displace it from the palmitoylation site (Stamler et al., 1992, Salaun et al., 2010). *S*-nitrosylation is involved in many intracellular signaling mechanisms. It is stimulus-evoked, precisely targeted, reversible, spatiotemporally restricted, and necessary for specific cellular responses (Hess et al., 2001, Hess et al., 2005, Seth et al., 2018). Independent proteomic studies have shown that both S-NO and S-PALM occur at numerous neuronal proteins with equal frequency and site specificity on the target molecule (Benhar et al., 2010, Doulias et al., 2010, Forrester et al., 2009, Hao et al., 2006, Jaffrey et al., 2001, Forrester et al., 2010, Percher et al., 2016, Wan et al., 2007). Ho et al. recently provided evidence of reciprocal regulation of the major postsynaptic protein-postsynaptic density protein 95 (PSD-95) by S-NO and S-PALM (Ho et al., 2011). They noted that the decrease in PSD-95 S-PALM increased its S-NO, suggesting that the specific sites of PSD-95 protein may undergo S-PALM or S-NO in a competitive manner. Interestingly, however, when S-palmitoylated PSD-95 localized in the synapse, it coordinated *N*-methyl-D-aspartate (NMDA) receptor stimulation and the subsequent production of nitric oxide by neuronal nitric oxide synthase. Nitric oxide, in turn, inhibited PSD-95 S-PALM and reduced the amount of this protein at the synapse.

A critical role for the crosstalk between S-PALM and S-NO in maintaining proper synaptic organization and function is further underscored by the widely reported imbalance in the dynamics of protein S-PALM and S-NO that are associated with the pathophysiology of neurodegenerative disorders, such as Alzheimer’s disease, Huntington’s disease, schizophrenia, mental disorders, and Parkinson’s disease (Akhtar et al., 2012, Barone et al., 2011, Butterfield and Sultana, 2007, Foster et al., 2003, Foster et al., 2009, Nakamura and Lipton, 2011, Zareba-Koziol et al., 2014). Still unknown, however, is whether disruption of the crosstalk between S-PALM and S-NO leads to the development of these diseases.

Considering recent insights into the role of rapid PTMs in the synaptic response to physiological stimuli and under pathological conditions, the present study investigated PTMs of endogenous synaptic proteins at the level of single cysteines. We employed a unique proteomic-based approach that integrates specific PSD protein enrichment, the chemical derivatization of modified sites and specific resin-based enrichment, label-free mass spectrometry (MS), and the differential analysis of two-dimensional (2D) heat-maps that represented the mass-to-charge ratio (m/z) *vs*. liquid chromatography (LC) retention times of each peptide ion (**Fig. 1**).

**Figure 1.**
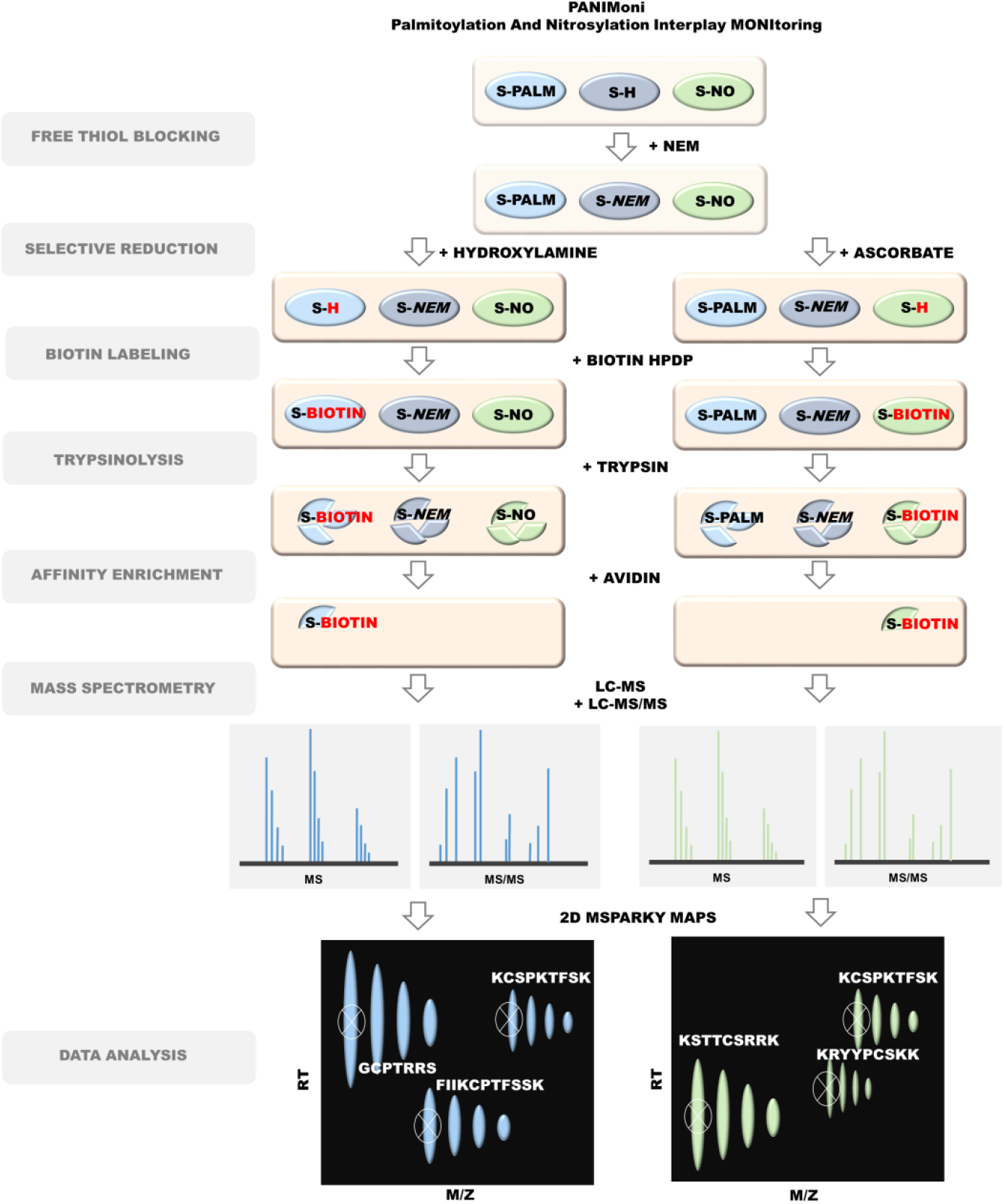
Schematic representation of PANIMoni proteomic approach for large-scale, site-specific S-palmitoylation and S-nitrosylation monitoring. Protein modifications were selectively replaced with biotin groups. After trypsin digestion and enrichment on avidin resin, the S-PALM and S-NO peptides were analyzed using MS in two independent runs (LC-MS and LC-MS/MS). In-house-developed MSparky software was used for site-specific data analysis.

We used this approach to assess dynamic, site-specific changes in two important cysteine modifications (S-PALM and S-NO) in a mouse model of chronic stress. We refer to this method as <u>Pa</u>lmitoylation And <u>N</u>itrosylation <u>I</u>nterplay <u>Moni</u>toring (PANIMoni).

Our results indicated that the mechanisms of crosstalk between the S-PALM and S-NO of proteins that are involved in synaptic transmission, synaptic localization, and the regulation of synaptic plasticity might be a major event that is associated with chronic stress-related disorders, leading to the rewiring of neuronal circuitry.

## Results

### The synaptic palmitoyl-proteome

To analyze the S-PALM of the synaptic proteome under physiological conditions, we used enriched fractions of PSD proteins that were obtained from brain synaptoneurosomes of control mice (**Fig. S1A**). Electron micrographs of synaptoneurosomes that were isolated by ultracentrifugation with a discontinuous Ficoll gradient revealed the well-preserved morphology of synaptic structures with a presynaptic mitochondrion, synaptic vesicles, and densely stained membranes that represented PSDs (**Fig. S1B**) (Zareba-Koziol et al., 2014). From the synaptoneurosomal fraction, we isolated enriched PSDs (350-450 µg/single brain) by sucrose density gradient centrifugation, which includes limited solubilization by the non-ionic surfactant Triton X-100. Using Western blot and MS, we confirmed that all of the fractions were enriched in PSD-95 protein, a postsynaptic marker, whereas the exclusively presynaptic protein synaptophysin was present only in synaptoneurosomal and homogenate samples (**Fig. S1C**). We identified a total of 3258 peptides that were assigned to 1128 PSD proteins. The intensities of the peptides that were identified in the fractions from two independent PSD preparations were compared on a scatter plot (**Fig. S1D**). The correlation coefficient between biological replicates was 0.9855, and the mean coefficient of variation (CV) was < 1%, indicating high reproducibility of the procedure. This optimized workflow for PSD isolation was used for all of the subsequent PTM analyses.

**Figure S1.**
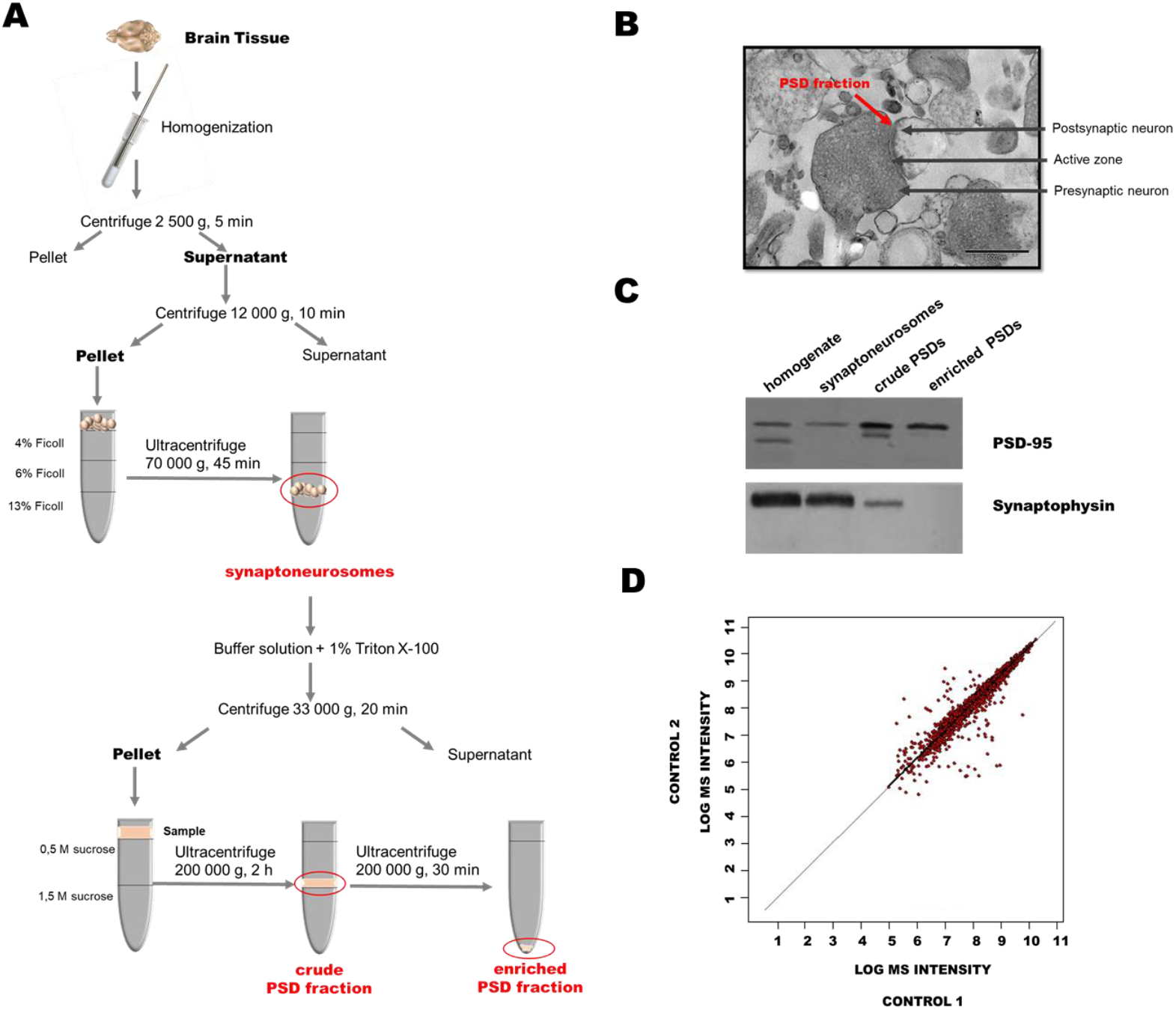
Workflow of PSD isolation and purity and reproducibility analysis. (A) Workflow of isolation of mouse brain postsynaptic densities (PSDs). (B) Electron micrographs of representative sample of synaptoneurosomes that were isolated from mouse brain. The arrow indicates components of the synapse: presynaptic terminal that contains vesicles and a postsynaptic element with PSD. (C) Representative Western blot analysis of postsynaptic and presynaptic markers in the homogenates, synaptoneurosomes, and PSDs that were isolated using optimized multistep ultracentrifugation. (D) Comparative analysis of postsynaptic density fraction preparation reproducibility that shows the peptide signal heights in two biological replicates of synaptosomal fractions. Each red dot corresponds to the intensity of the same peptide that was identified in the first (vertical axis) and second (horizontal axis) replicates. A total of 3258 peptides were included in the analysis. The mean coefficient of variation was less than 1%.

Because of an insufficient amount of protein for PTM analysis that could be obtained from one mouse brain, each sample contained PSDs from two freshly isolated mouse brains. To assess whether the PSD enrichment protocol enabled the identification of endogenously *S*-palmitoylated proteins, we used the Acyl-Biotin Exchange (ABE) assay, based on the labeling of S-PALM proteins with a biotin moiety at the site of a modified cysteine. Biotinylated proteins were visualized by immunoblotting with an anti-biotin antibody (**Fig. 2A**). This analysis revealed numerous protein bands across a broad mass range in the PSD fractions that were isolated from the brains of control mice, indicating the presence of endogenously S-PALM proteins. The selectivity of ABE was confirmed by the lack of a signal from biotinylated proteins in samples that were not treated with hydroxylamine, which reduced *S*-acyl bonds (**Fig. 2B**).

**Figure 2.**
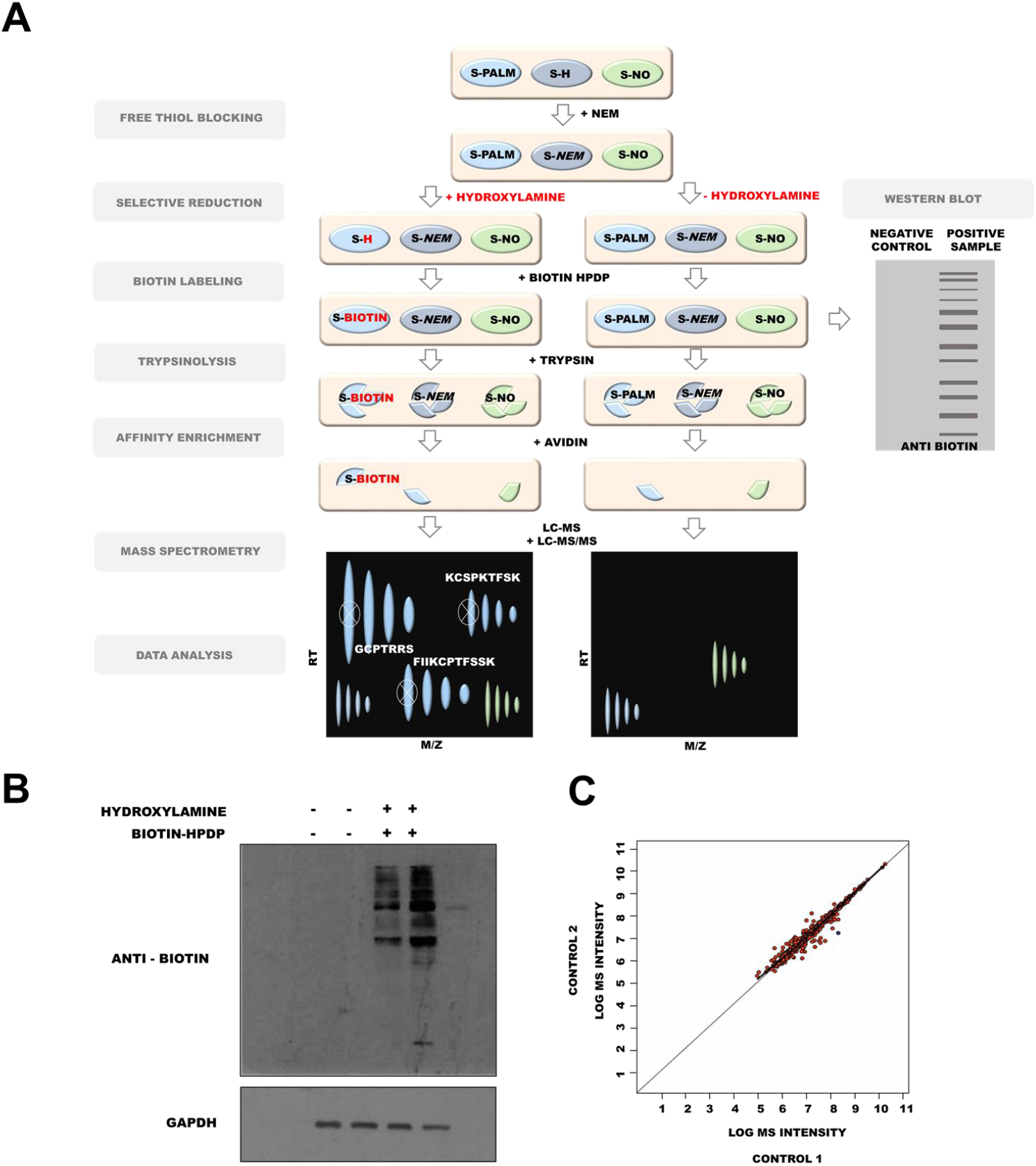
S-palmitoylome of mouse PSD proteins. (A) Schematic representation of S-palmitoylation analysis. (B) Western blot analysis of S-palmitoylation in the PSD protein fraction. (C) Mass spectrometry-based analysis of S-palmitoylation that shows a high correlation between biological replicates (CV < 1%).

We then sought to identify the exact sites of *S*-palmitoylated cysteines. To analyze the S-palmitoylome of the PSD fractions, we used a high-throughput proteomic PANIMoni approach. This approach utilizes the PalmPISC (palmitoyl protein identification and site characterization) method [65] to simultaneously identify S-PALM sites and their cognate proteins in complex biological mixtures, combined with the analysis of 2D heat-maps that represent the *m/z vs*. LC retention times of each peptide ion. Importantly, S-PALM proteins were distinguished from nonspecifically enriched proteins by comparing samples that were treated in parallel with or without hydroxylamine (**Fig. 2A**). Using MSparky software, 2D heat-maps were generated, and the peptide signals on the 2D heat-maps that were obtained for different biological samples were correlated and labeled with appropriate peptide sequences. The merged list of all S-PALM PSD peptide sequences that were acquired in the entire LC-MS and LC-MS/MS experiments was used for signal assignments in a 2D heat-map for a single biological replicate. Nonspecifically enriched proteins were excluded based on a simultaneous analysis of the hydroxylamine-free negative control.

Using the PANIMoni approach, 705 S-PALM cysteine sites were identified on 404 distinct proteins in PSD fractions that were isolated from the mouse brains. In all of the samples, we identified mainly cysteine-containing peptides. The MSparky 2D heat-map analysis showed a high correlation between biological replicates (CV < 1%, *R* = 0.98), indicating the high reproducibility of S-PALM peptide enrichment and measurement (**Fig. 2C**). The same sites of S-PALM were found in three independent biological replicates. Cysteine-containing peptides and assigned proteins that were identified with high stringency in the MS/MS experiments from three biological replicates are listed in **Table S1**. Among them, 252 S-PALM proteins have been previously shown to undergo S-PALM or to have their activity altered by this modification, whereas 152 proteins (from 404 listed) represented newly identified targets of S-PALM.

### Dynamic changes in S-palmitoylation in an animal model of stress-related diseases

To determine whether the changes in protein S-PALM are associated with the chronic stress response, we used a mouse model of chronic restraint stress in which aberrant plasticity has been well documented (**Fig. 3A**) (Conrad et al., 2004, Conrad, 2008, McLaughlin et al., 2005, McLaughlin et al., 2007, Wright and Conrad, 2005, Duman and Aghajanian, 2012). Mice that were subjected to restraint stress for 21 days (6 h/day) exhibited depressive-like behavior (i.e., an increase in immobility time in the tail suspension test, *p* < 0.05), elevated serum corticosterone levels (*p* < 0.01), and a decrease in body mass compared with control animals (**Fig. 3B-D**).

**Figure 3.**
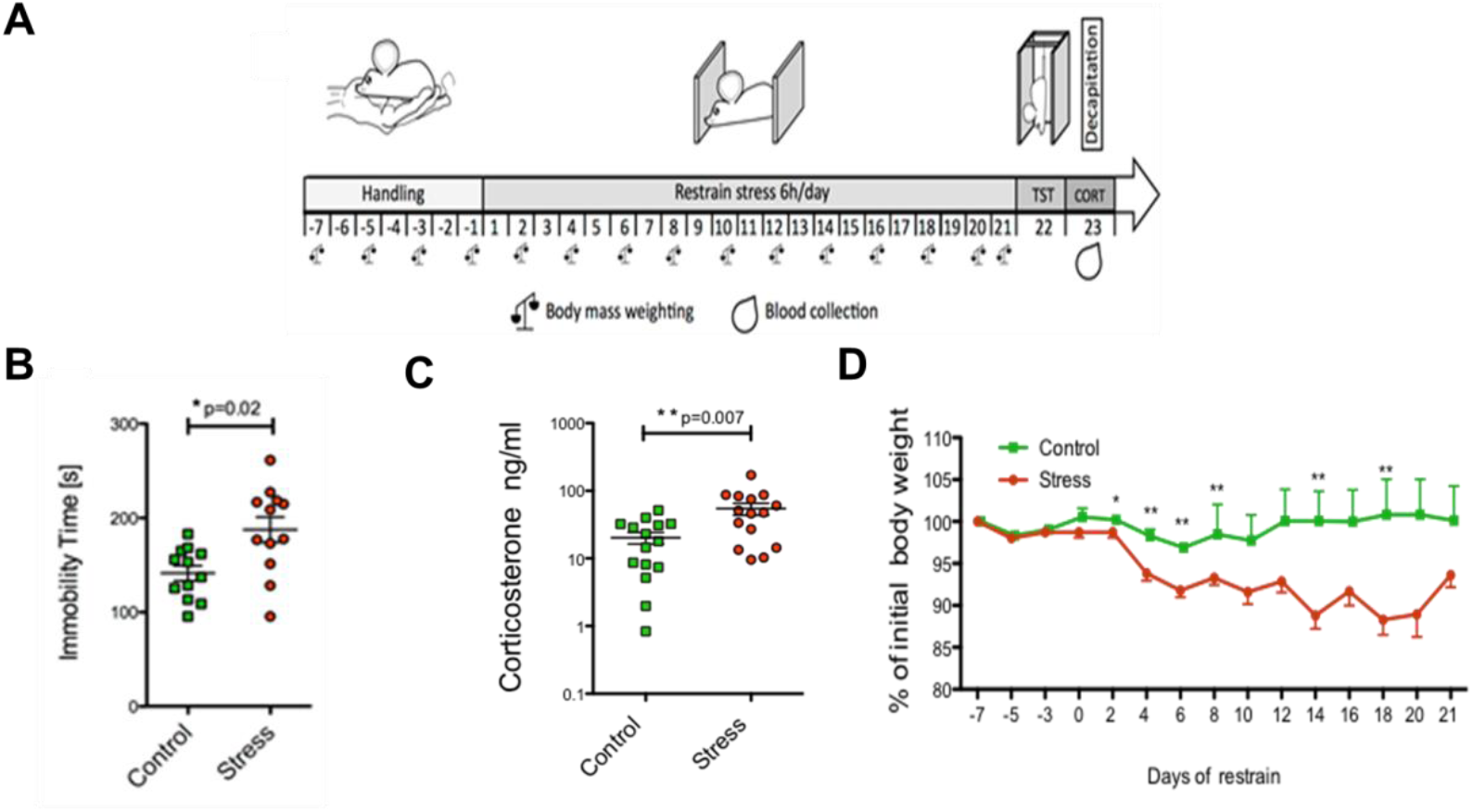
Schematic representation of the mouse model of chronic restraint stress and behavioral tests. (A) Schematic illustration of mouse treatment. (B) Tail suspension test. (C) Corticosterone levels. (D) Changes in body mass recorded every other day throughout the experiment.

Because changes in protein expression may interfere with the palmitoylome profile, we first analyzed the effects of chronic restraint stress on the level of PSD proteins. By employing a quantitative proteomics approach (label-free method coupled with 2D heat-map analysis) we identified 1247 protein clusters (single proteins or clusters of gene identifiers that matched the same set of peptides) that were represented by at least two peptides and met other inclusion criteria, such as the correct *m/z*, retention time, and peak envelope (**Table S2**).

Principal component analysis (PCA) was used to assess the level of similarity of protein content. Within each group of control mice (*n* = 3) and chronically stressed mice (*n* = 3), we confirmed the reproducibility of the PSD proteome preparation but also found a clear distinction between control and stressed mice (**Fig. 4A**). The results of the relative quantification experiment are summarized in **Fig. 4B**. The raw protein ratios and *q* values are listed in **Table S1**. We identified six significantly changed proteins: four upregulated proteins with a fold change > 1.25 and *q* < 0.05 (protein SZT2, LIM zinc-binding domain-containing Nebulette, probable E3 ubiquitin-protein ligase MID2, 60S ribosomal protein L11) and two downregulated proteins with a fold change < 1.25 and *q* < 0.05 (V-type proton adenosine triphosphatase [ATPase] 16 kDa proteolipid subunit and Leucine zipper putative tumor suppressor).

**Figure 4.**
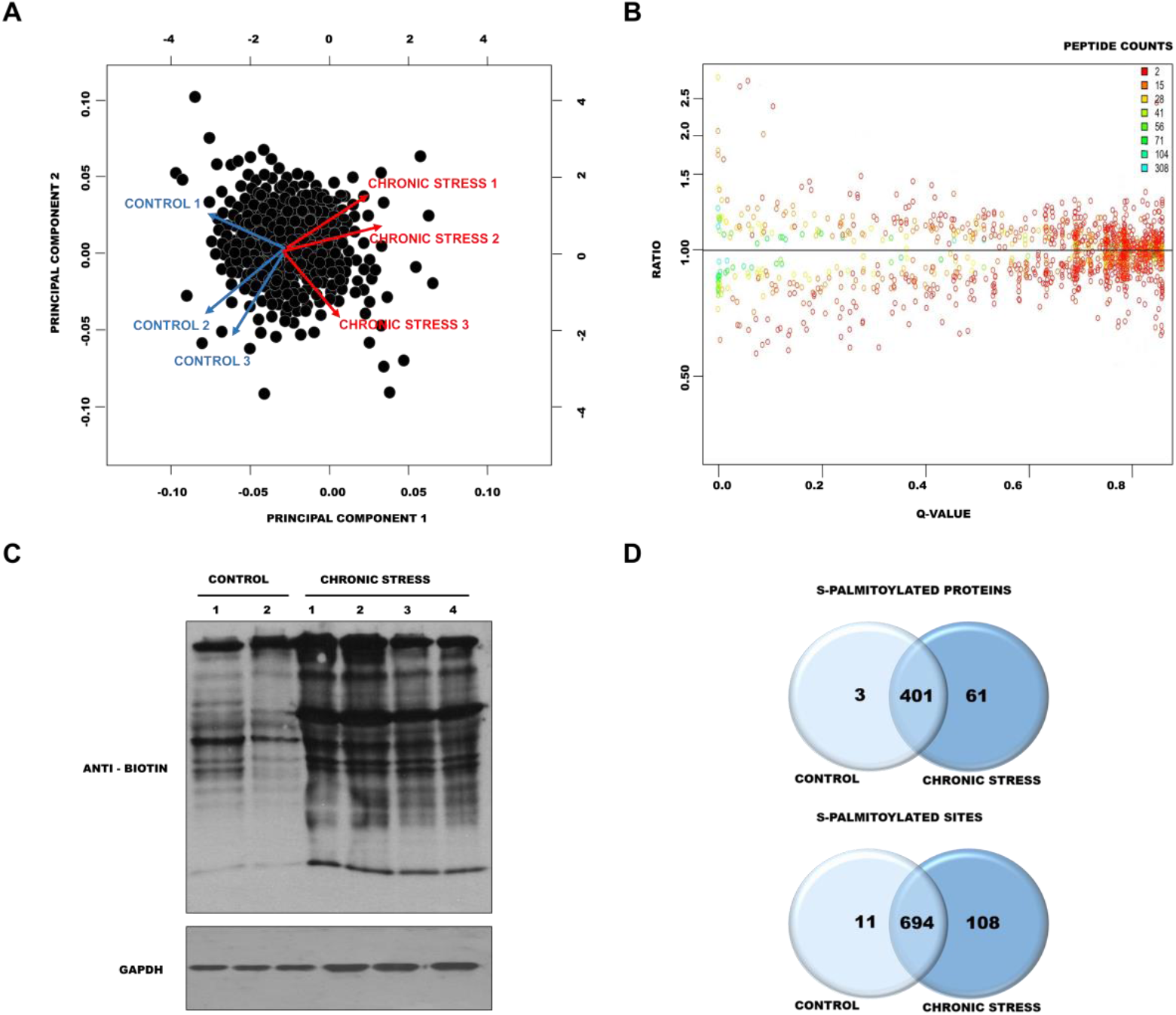
Analysis of changes in protein expression and S-palmitoylation in the brains of mice after chronic restraint stress. (A) Principal Component Analysis of identified proteins in control and chronically stressed mice. (B) Distributions of the log-ratios (vertical axis) and q values (horizontal axis) of proteins from control mice and chronically stressed mice (data analyzed with Diffprot). (C) Western blot analysis of S-palmitoylation pattern in control mice and chronically stressed mice. S-palmitoylation sites were selectively labeled with S-S-biotin (ABE). The visualization of biotinylated proteins was achieved using anti-biotin antibodies. (D) Venn diagram comparisons of the numbers of S-PALM proteins and sites that were identified in synaptic protein fractions from control mice and chronically stressed mice.

Next, to determine whether chronic stress influences protein S-PALM, we analyzed changes in the patterns of this modification. Using the ABE method combined with Western blot, we observed an increase in the number and intensity of bands that represented S-palmitoylated proteins in PSDs from stressed mice (**Fig. 4C**, lanes 3-6) compared with control mice (**Fig. 4C**, lanes 1 and 2). To further confirm that stress increases the S-PALM of proteins, we performed a detailed, differential analysis of S-PALM at the level of single cysteines using the PANIMoni approach. We identified 813 S-PALM cysteine sites that were assigned to 465 proteins in the PSD fractions (**Table S3A-C)**. Among the 465 S-PALM PSD proteins that were identified, three were found only in control mice, and 61 were exclusively found in stressed mice (**Fig. 4D**). We also identified proteins that differed at the level of modified sites. A total of 813 sites were identified, including 108 sites that were found exclusively in PSDs in stressed mice and 11 that were uniquely detected in control PSD fractions.

### Mechanism of dynamic S-palmitoylation regulation

Cysteines of a single protein may be either *S*-palmitoylated or *S*-nitrosylated, depending on the physiological context. This may represent a general mechanism of the modulation and fine-tuning of molecular signaling, but it has not yet been characterized in a global type of analysis. To explore this possibility, we analyzed changes in S-PALM in primary mouse cortical neurons that were treated with the stress hormone corticosterone. Western blot and *in situ* cell imaging revealed that corticosterone increased the level of *S*-palmitoylated proteins in cultured neurons (**Fig. 5A panel 1**, **Fig. 5B lane 1**), which corresponded to our *in vivo* results. Interestingly, treatment of the cultures with the nitric oxide donor *S*-nitroso-*N*-acetyl-DL-penicillamine (SNAP) or S-PALM inhibitor 2-bromopalmitate decreased the level of global S-PALM, whereas the nitric oxide synthase inhibitor *N*_ω_-nitro-L-arginine methyl ester hydrochloride (L-NAME) caused an opposite effect, increasing protein palmitoylation (**Fig. 5A panel 2-5**, **Fig. 5B lane 2-5**). Elevated S-PALM is thus an element of the neuronal response to high corticosterone levels and can be regulated by nitric oxide.

**Figure 5.**
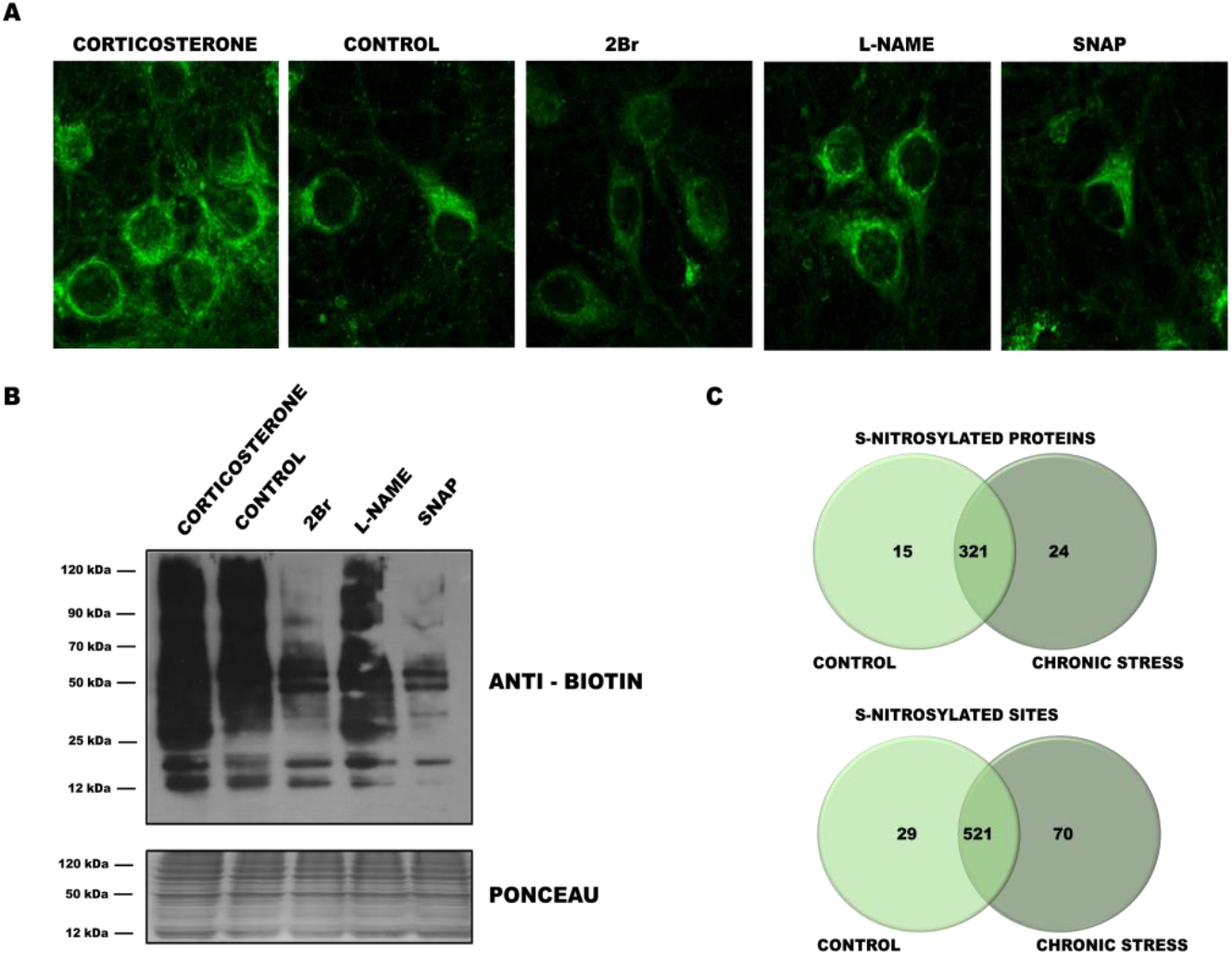
Crosstalk between protein S-palmitoylation and S-nitrosylation. (A) Imaging of total proteome palmitoylation in neuronal cultures subjected to click chemistry reaction with Oregon Green 488 dye. (B) Western blot ABE analysis applied to detection of the total S-palmitoylome in primary cortical neurons that were treated with corticosterone, 2Br, SNAP, and L-NAME. (C) Venn diagram comparisons of the numbers of S-NO proteins and sites that were identified in synaptic protein fractions from control mice and chronically stressed mice.

To further characterize crosstalk between S-PALM and S-NO, we analyzed the global protein S-NO profile in the chronic restraint stress model of aberrant synaptic plasticity in mice using the PANIMoni approach. The analysis generated lists of peptides that were obtained separately for stressed mice and age-matched controls. Protein IDs were assigned to the appropriate peptide sequences and are listed in **Table S4A-C**.

A total of 620 S-NO cysteine sites that were assigned to 360 proteins were identified among PSD proteins (**Fig. 5C**). A total of 39 differential protein species were identified. Fifteen proteins appeared only in controls, whereas 24 proteins were modified only in PSD fractions that were obtained from stressed mice. Of the 620 S-NO sites that were identified, 29 were found only in control PSDs, and 70 were detected only in stressed mice. These nitrosopeptides represented 63 uniquely regulated proteins after chronic stress (**Table S4**).

Complete lists of *S*-palmitoylated and *S*-nitrosylated PSD proteins from the brains of chronically stressed and control mice enabled us to perform detailed comparisons of these PTM profiles at the level of single cysteines. The results of the high-throughput MS experiments revealed two sets of S-PALM proteins and two sets of S-NO proteins that corresponded to the chronic stress model and age-matched controls.

Of the 446 modified PSD proteins (778 cysteines) that were identified in control mice, 42 were exclusively *S*-nitrosylated, and 109 were exclusively *S*-palmitoylated (**Fig. 6A**). Our data revealed that the majority of the PSD protein cysteines that were detected (58% of the proteins that were identified) occurred only in one of the PTM types in control mice.

Chronic stress generally results in a greater number of modified cysteine sites that undergo both S-PALM and S-NO. Interestingly, in chronically stressed mice, among the 464 modified proteins that were identified, only two were exclusively associated with nitric oxide, and 119 were exclusively associated with palmitate (**Fig. 6B**). Notably, the differences between the sets of S-NO and S-PALM proteins at the cysteine level were also much less pronounced, and only seven peptides were exclusively modified by nitric oxide. The remainder of the *S*-nitrosylated sites and proteins that were identified existed in two modified forms after chronic stress. The crosstalk between the S-NO and S-PALM of brain PSD proteins appears to vary, depending on the physiological condition.

**Figure 6.**
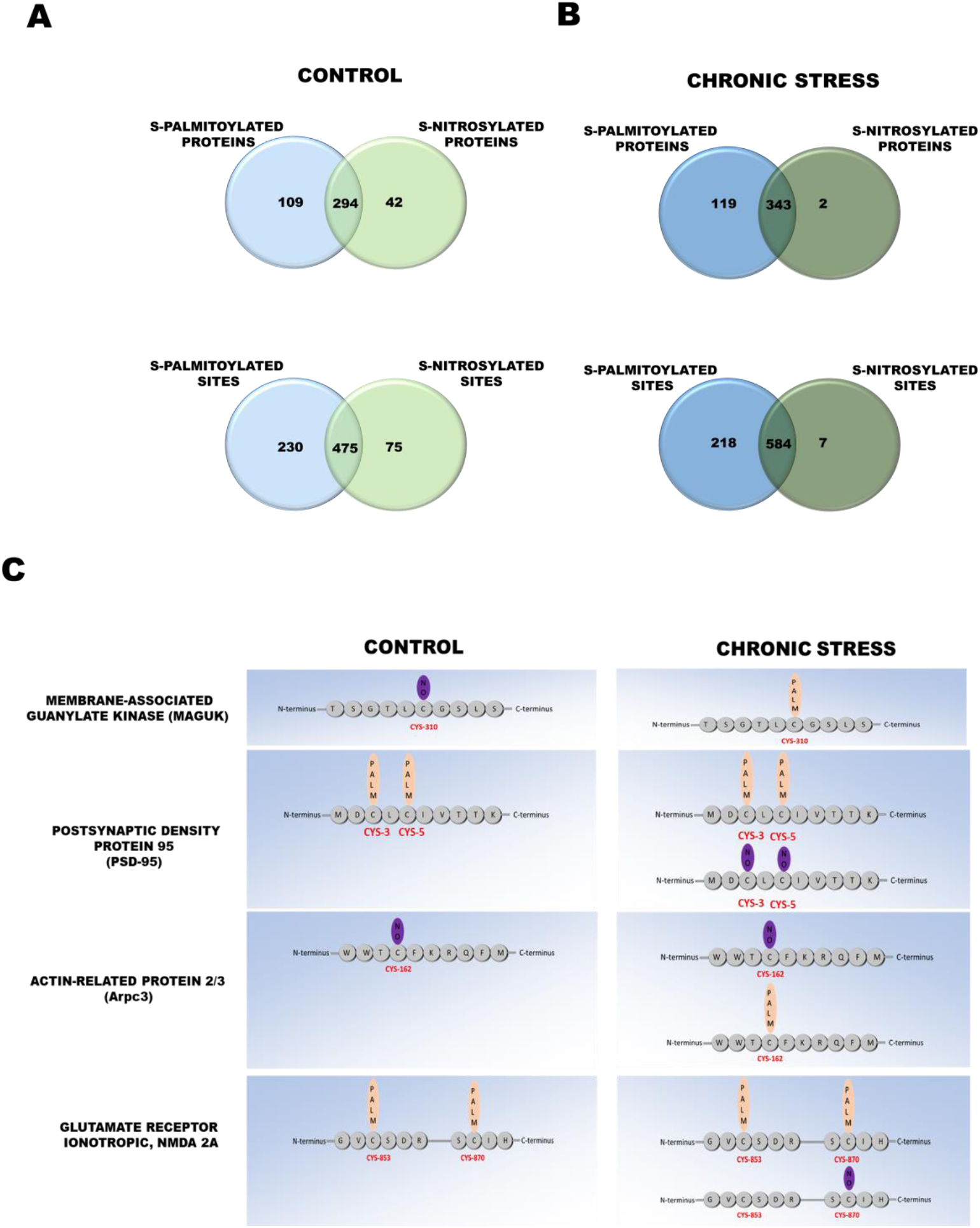
Aberrant S-palmitoylation/S-nitrosylation crosstalk caused by chronic stress. (A) Venn diagram comparisons of the numbers of S-PALM and S-NO proteins and sites that were identified in synaptic protein fractions from control mice (B) and chronically stressed mice. (C) S-PALM pattern in control mice and chronically stressed mice that shows differential S-PALM for important synaptic proteins.

Finally, among a total of 813 S-PALM and 620 S-NO cysteine sites that were characterized on 465 and 360 proteins, respectively, we sought to identify those that were differentially affected by stress. We obtained evidence that 443 cysteine sites (54% of all of the sites that were identified) that belonged to 297 proteins in all of the experimental conditions occurred in both S-PALM and S-NO forms. Interestingly, the remainder of the cysteines that were identified (46%) underwent regulation that strictly depended on the biological conditions.

A total of 123 of all of the modified proteins underwent changes in the patterns of S-NO and S-PALM that were induced by chronic stress. All this protein grouped according to their molecular function are presented in **Table 1**. Surprisingly, for some of the proteins, such as Iqsec2 and Maguk, we observed an exchange of S-PALM to S-NO after chronic stress. All types of molecular switches that were found between these two PTMs that were triggered by chronic stress are shown in **Fig. 6C**.

**Table 1.** Identified PSDs proteins with affected S-PALM/S-NO crosstalk

| **Protein class** | **Gene Name** | **Palmitoyl-proteome studies** |
| --- | --- | --- |
| Cell adhesion proteins | Ctnna1(1), Cntnap1(2), Ncam1(3), Ctnnd1(4), Ctnnb1(5), Jup(6), Jam3(7), Ctnnd2(8), Src(9) | Shen LF et al. 2017(1,5), Collins MO et al. 2017(1,4,5,8), Chesarino NM et al. 2014 (1,3,4), Wan J et al. 2013(2,3, 8) Ren W et al. 2013(2,3,4), Sobocińska J et al. 2018(4), Shen LF et al. 2017(4,6,8), Gould NS et al. 2015(4,5,6), Li Y et al.2012(4,7) Lievens PM et al. 2016(3), Ponimaskin E et al. 2008(3), Stypulkowski E et al. 2018(4), Roberts BJ et al. 2014(6) |
| Cytoskeletal proteins | Tubb4a(1), Map1a(2), Ctnna1(3), Ablim1(4), Arpc3(5), Map1b(6), Dync1h1(7), Ctnnd1(8), Tuba4a(9), Plec(10), Tuba1a(11), Fscn1(12), Myo5a(13), Coro1c(14), Sept4(15), Opa1(16), Ctnnd2(17), Epb41l3(18), Inf2(19), Map6d1(20), Iqsec2(21), Ank2(22), Iqsec1(23), Actn1(24), Ank3(25), Sptbn1(26), Syne1(27), Cyfip1(28), Phactr1(29) | Ren W et al. 2013(1,8,10,12,26), Collins MO et al. 2017(2,3,8,17,22), Shen LF et al. 2017(3,8,9,10,24), Chesarino NM et al. 2014 (3,8,12,5,10,26,28), Wan J et al. 2013(6,12,13,18,22,25,26,28), Sobocińska J et al. 2018(8), Gould NS et al. 2015(8,24,26), Li Y et al.2012(8), Kang R et al. 2008(14,16), Wei X et al. 2014(19), Thinon E et al. 2016(24,28)<br>Kang R et al. 2008 (17), Gory-Fauré S et al. 2006(20), |
| Structural proteins | Crocc(1), Ina(2), Bsn(3), Pclo(4) | Pinner AL et al. 2016(1), Hernandez JL et al. 2016(1), Wan J et al. 2013(2,3), Collins MO et al. 2017(3) |
| Chaperonins | Ywhah(1), Dnajc11(2) | Sobocińska J et al. 2018, Chesarino NM et al. 2014, Shen LF et al. 2017(2), ), Collins MO et al. 2017(2), Ren W et al. 2013(2), Martin BR et al. 2011(2) |
| Cytokine | Cmtm4 |  |
| Receptors/channels | Adgrb2(1), Leprotl1(2), Gabbr2(3), Gria1(4), Atp1a3(5), Adam22(6) | Shen LF et al. 2017(2), Collins MO et al. 2017(2,5), Pinner AL et al. 2016(4), Wan J et al. 2013(5)<br>Spinelli M et al. 2017(4), Hayashi T et al. 2005 (4) |
| Ion/electron transporting proteins | Ndufs2(1), Ndufs1(2), Dld(3), Slc25a1(4), Slc25a3(5), Slc12a5(6) | Shen LF et al. 2017(1,2,3,5), Ren W et al. 2013(1,2,3,4,5), Collins MO et al. 2017(2,5), Gould NS et al. 2015(2,3,4), Wan J et al. 2013(2,3,5,6), Thinon E et al. 2016(3,5), Sobocińska J et al. 2018(5), Chesarino NM et al. 2014(5) |
| Receptor binding proteins/receptor modulators | Lgi1, Ctnnb1(2), Prkcb(3), Ly6h(4), Ank2(5) | Kang R et al. 2008(1), Shen LF et al. 2017(2), Collins MO et al. 2017(2,3,5), Gould NS et al. 2015(2), Wan J et al. 2013(3,5), Stypulkowski E et al. 2018(2) |
| Transferases | Mecp2(1), Acly(2), Crat(3), Tst(4), Acadsb(5), Gsk3b(6), Cdc42bpb(7), Comtd1(8), Prkcb(9), Mpp2(10), Cs(11), Gk(12), Got2(13), Src(14), Hk1(15), Pkm(16), Ogt(17), Kalrn(18), Begain(19), Ngef(20), <u>Tst</u>(21) | Sobocińska J et al. 2018(2,13,15), Shen LF et al. 2017(2,4,5,13,16), Thimon E et al. 2016(2,13,15), Gould NS et al. 2015(2,4,12,13), Chesarino NM et al. 2014(2,13), Wan J et al. 2013(2,9,10,13,15,16), Collins MO et al. 2017(9,10,15), Edmonds MJ et al. 2017(11), Yount JS et al. 2010(13), Ren W et al. 2013(15,16), Li Y et al. 2012 (15), Dowal L et al. 2011(18) |
| Protein transporters | Dync1h1(1), Ap1b1(2), Ap3d1(3), Exoc1(4), Exoc2(5), Exoc5(6), Mtx1(7), Ank2(8), Snph(9) | Shen LF et al. 2017(7), Gould NS et al. 2015(1,2), Collins MO et al. 2017(1,8,9), Wan J et al. 2013 (1,2,8),Chesarino NM et al. 2014(1), Thimon E et al. 2016(3) |
| GTPases/signaling proteins | Homer1(1), Kalrn(2), Plcb1(3), Sirpa(4), Rasal1(5), Tubb4A(6), Gnai2(7), Eef1a2(8), Tbc1d24(9), Kras(10), Sept4(11), Opa1(12), Rap1b(13) | Edmonds MJ et al. 2017(1), Collins MO et al. 2017(2,3,5,7,8), Wan J et al. 2013(2,3,7,8,10), Ren W et al. 2013(6,7,8), Sobocińska J et al.(7), Shen LF et al. 2017(7), Thimon E et al. 2016(7,13), Chesarino NM et al. 2014(7,10,13 ), Li Y et al. 2012(7), Merrick BA et al. 2011(7,10), Martin BR et al. 2011(7,13), Yount JS et al. 2010(7), Kang R et al. 2008(12), |
| GTPase modulating proteins | Rasa3, Sbf1(2), Ngef(3) |  |
| Nucleic acid binding proteins | Rps3a(1), Sirt2(2), Eef1a2(3), Mecp2(4), Tst(5), | Shen LF et al. 2017 (1,5), Gould NS et al. 2015(1,5), Chesarino NM et al. 2014 (1), Collins MO et al. 2017(2,3), Wan J et al. 2013(3), Ren W et al. 2013(1,3), Yount JS et al. 2010 (1) |
| Enzymatic activity proteins | Plcb1(1), Prkcb-2, <u>Src</u>-3, Hk1-4, Adam22-5, Bphl-6, Ass1-7, Acly-8, Aldoa-9, Idh3a-10, Pdha1-11, Mdh1-12, Acadsb-13, Ogdh-14, Aldh3b1-15, Marc2-16, Pdhb-17, Got2-18, | Collins MO et al. 2017(1,4,11,14), Wan J et al. 2013(1,4,8,10,11,12,14,17,18), Sobocińska J et al. 2018(4,7,8,10,12,14,15,18), Thimon E et al. 2016(4,8,10,12), Ren W et al. 2013(4,10,11,12,14,15,17), Li Y et al. 2012(4), Shen LF et al. 2017(6,7,8,11,12,13,14,16,17,18) , Gould NS et al. 2015(6,7,8,10,11,12,16,18), Chesarino NM et al. 2014(6,8,10,12,14,15,17,18), Yount JS et al. 2010 (18) |
| Scaffolding Synaptic proteins/ Synapse organization | Pclo, Homer-2, Mpp2-3, Shank2-4, Dlgap2-5, Ank3-6, Lrrtm3-7, Bsn-8 | Edmonds MJ et al. 2017(2), Collins MO et al. 2017(3,5,8), Wan J et al. 2013(3,6,8) |
| Developmental proteins | Ctnnd2-1, Snph-2, Cyfip1-3, Hapln1-4, Ngef-5, Ina-6, Dip2a-7, Tspan2-8, | Shen LF et al. 2017(1), Collins MO et al. 2017(1,2,8), Wan J et al. 2013(1,3,6), Thinon E et al. 2016(3), Chesarino NM et al. 2014(3) |
| Other | Tmem50b-1, Wdr7-2, Strn4-3, Ajm1-4, Snph (5), Dip2b-6, Amer-7, Nipsnap2-8, Ncdn-9 | Collins MO et al. 2017(1,5,8,9), Wan J et al. 2013(2,9), Thinon E et al. 2016(7), |

### S-palmitoylation and S-nitrosylation affect the function of various PSD proteins

To investigate the biological importance and functional significance of PSD proteins that exhibited changes in S-PALM/S-NO crosstalk, we performed functional enrichment analysis using Gene Ontology (GO) and the Kyoto Encyclopedia of Genes and Genomes (KEGG) (Kanehisa and Goto, 2000, Mi et al., 2017). Overrepresentation statistics were used to calculate the probability of highly populated protein classes, pathways, and GO terms among S-PALM and S-NO proteins that occurred in a non-random manner. Indeed, many of the protein categories were overrepresented in the data. As a reference, we used the set that comprised 11970 proteins that were associated with PSDs. The reference set was generated from our experimental data and the Genes to Cognition database (Croning et al., 2009, Grant et al., 2005).

Gene Ontology biological process (GO_BP) analysis linked a majority of the proteins with stress-induced changes in S-PALM/S-NO crosstalk to processes that are associated with synaptic plasticity, such as energy metabolism (fold change = 27.93, *p* = 5.17E-06, tricarboxylic acid metabolic process; fold change = 8.12, *p* = 4.91E-06, generation of precursor metabolites and energy), acylation metabolism (fold change = 23.1, *p* = 2.32E-02, acetyl-CoA metabolic process; fold change = 12.66, *p* = 1.24E-02, thioester metabolic process; fold change = 9.91, *p* = 2.98E-03, oxidoreduction coenzyme metabolic process), and protein localization (fold change = 12.66, *p* = 1.24E-02, actin cytoskeleton organization; fold change = 12.66 *p* = 1.24E-02, actin filament-based process; **Fig. 7A**). Furthermore, proteins that are related to synaptic dynamics, the positive regulation of synaptic transmission, the regulation of synaptic plasticity, and the modulation of synaptic transmission were also significantly overrepresented.

**Figure 7.**
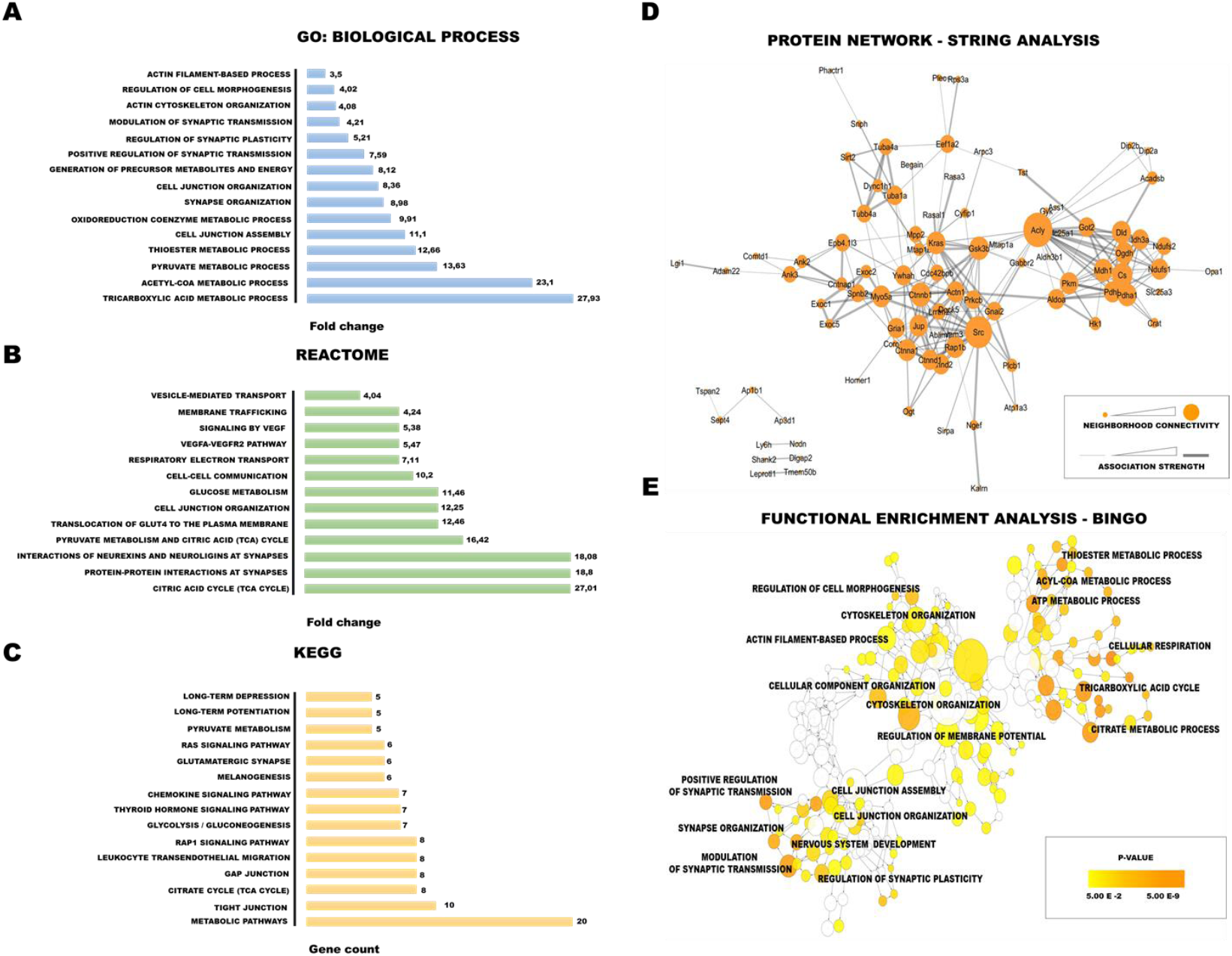
Bioinformatics tools for the analysis of proteins with aberrant S-PALM/S-NO crosstalk. (A-C) Gene Ontology analysis in terms of (A) biological processes (GO_BP), (B) REACTOME pathway analysis, (C) KEGG pathway analysis (D) STRING analysis of protein interactome. (E) Bingo analysis of enriched biological classes.

We also visualized biological process overrepresentation using the BINGO plugin (**Fig. 7B**). To visualize GO terms and biological pathways that are associated with synaptic proteins that exhibited stress-induced changes, we used BINGO bioinformatic software in the integrative environment of Cytoscape 3.1. Significantly overrepresented biological processes, based on GO terms, were visualized in Cytoscape. The size of a node is proportional to the number of targets in the GO category. The color represents enrichment significance. A deeper color on the color scale indicates higher enrichment significance. *p* values were adjusted using Benjamini and Hochberg False Discovery Rate (FDR) correction.

We also analyzed proteins that exhibited changes in PTM crosstalk using REACTOME pathways. Most of the significantly enriched (*q* < 0.05) proteins are related to energy metabolism (e.g., glucose metabolism, TCA cycle, and pyruvate metabolism), protein localization (e.g., membrane trafficking and the translocation of GLUT4 to the plasma membrane), and synapse signaling (e.g., signaling by vascular endothelial growth factor, interactions of neurexins and neuroligins at synapses, and protein-protein interactions at synapses; **Fig. 7C**).

The KEGG pathway enrichment results associated the target proteins with both energy metabolism (citrate cycle [TCA cycle], glycolysis/gluconeogenesis, and pyruvate metabolism) and synaptic plasticity (long-term depression, long-term potentiation, and glutamatergic synapse-connected terms as enriched), which was consistent with the GO and KEGG analyses (**Fig. 7D**).

Most proteins exert their biological functions by interacting with each other. To uncover functional aspects that are associated with proteins that are affected by chronic stress, we used the STRING database. Overall, at a medium STRING confidence level > 0.7, 84.0% of the of identified proteins were grouped in a single cluster, with an enrichment ratio of 3.15 over the randomly expected ratio. Furthermore, two smaller connection clusters were identified (**Fig. 7E**). Likewise, proteins that had changes in S-NO/S-PALM balance were associated with synaptic plasticity regulation, protein localization, and energy metabolism.

Our dissected analysis uncovered a remarkable degree of the stress-induced disruption of S-PALM/S-NO crosstalk in proteins that play key roles in proper synaptic function. This phenomenon may be a major event that is involved in the development of stress-related aberrant plasticity.

## Discussion

Crosstalk between different PTMs of proteins is an emerging theme in neurobiology. Proteins can be modified by multiple PTMs that acting as specific PTM code that modulate protein function. Proteins are frequently modified not only by a single type of PTM but also by several different PTMs. Therefore, to elucidate the role of individual PTMs in the context of specific protein activities, we must first understand their co-occurrence in the same protein or at the same protein site and their functional profile. These reciprocal dynamics between S-PALM and S-NO have been described in the context of synaptic proteins (Ho et al., 2011). These studies, however, were limited to single proteins and did not consider the global effects of alterations of S-PALM/S-NO crosstalk in complex physiological states.

For the purpose of the present study, we designed and developed a new proteomic approach called PANIMoni. It is an MS-based method that allows the differential, site-specific identification of PTMs of endogenous synaptic proteins. PANIMoni, in contrast to commonly used spectral counting methods, relies on peptide intensity quantification combined with protein identification. Its high sensitivity makes it suitable for comprehensively assessing dynamic changes in protein PTMs under different physiological and pathological conditions. Furthermore, our approach can be extended to quantitatively analyze different types of PTMs, such as sulfenylation and glutathionylation, in which the selective liberation of modifications and switch to biotin are possible.

Using the PANIMoni approach, we were able to perform a large-scale analysis of cysteine PTMs, namely S-PALM and S-NO, within PSD proteins of the mouse brain. Our goal was to test the hypothesis that stress-related aberrant plasticity is associated with global changes in the reciprocal interplay between these two PTMs. By reducing the complexity of the system that was analyzed and narrowing our research to analyzing PSD proteins in a well-defined mouse model of chronic stress, we obtained highly reproducible and informative results. The high-throughput analysis showed that postsynaptic proteins are strictly regulated by both cysteine modifications, S-PALM and S-NO, under physiological conditions. Our detailed analysis strictly defined 813 *S*-palmitoylated sites that were assigned to 465 proteins and 620 *S*-nitrosylated sites that were assigned to 365 proteins.

Importantly, the majority of the proteins were exclusively modified with one of the two PTMs. A competitive relationship between S-PALM and S-NO for neuronal proteins, including SNAP-25 and PSD-95, has been previously reported (Ho et al., 2011). Similar to previous studies, the proteins that we identified were associated with various signaling pathways (e.g., axon guidance, synaptic plasticity, and protein localization) and cell energy metabolism.

We also provide insights into the ways in which chronic stress affects S-PALM and S-NO. The analysis of PSD fractions from the brains of chronically stressed mice showed a substantial increase in the overall number of palmitoylated proteins compared with control animals. Interestingly, this elevation of protein S-PALM correlated with changes in the number of both nitrosylated cysteines and proteins.

In summary, we defined a very stringent set of 123 proteins that exhibited alterations of S-PALM/S-NO crosstalk after chronic stress at the level of specific sites of modification. Using functional bioinformatics analysis with various tools (i.e., REACTOME, KEGG, and GO), we assigned these 123 proteins to molecular pathways that are involved in synaptic transmission, protein localization, and energy metabolism. Notably, among this group of proteins, catenin delta CaMKII, Iqseq7, Shank, Gria1, glycogen synthase kinase-3β (GSK3β), and PSD-95 play key roles in regulating synaptic plasticity. Synaptic plasticity is involved in various stress-related disorders, and changes in protein PTM profiles may be a particularly important mechanism in the development of neuropsychiatric disorders.

GSK3β activity is altered by chronic stress and in schizophrenia and depression (Jope and Roh, 2006, Li and Jope, 2010). Previous studies have reported the S-NO-mediated activation of GSK3β (Ryu et al., 2016). The present data suggest that GSK3β activity may also be modulated by S-PALM, and S-PALM/S-NO crosstalk is crucial for GSK3β activity. We also found that chronic stress influenced the S-PALM/S-NO crosstalk of several neuronal kinases, including CaMKII and protein kinase C (PKC). CaMKII and PKC are activated by calcium influx in the synapse, resulting in an increase in synaptic strength (Lisman et al., 2012, Chu et al., 2014). The activity of these kinases is involved in actin polymerization, membrane trafficking, and glutamate receptor trafficking. CaMKII is upregulated in numerous diseases, including stress-related disorders, and stress hormones have been shown to significantly alter their functions (Sanhueza et al., 2011, Ghosh and Giese, 2015). *S*-nitrosylation-dependent CaMKII regulation has been recently reported. The present study suggests a possible link between alterations of the S-PALM/S-NO crosstalk of CaMKII and the response to chronic stress (Erickson et al., 2015). Similar to CaMKII, changes in PKC activity have been associated with behavioral effects of chronic stress, and these effects were mediated by the formation of nitric oxide, which is a substrate for S-NO (Chen et al., 2009). Alterations of PKC activity following chronic stress may thus be regulated by crosstalk between two cysteine modifications (S-PALM and S-NO).

In the present study, cytoskeletal proteins were abnormally modified by chronic stress. The dynamics of the synaptic cytoskeleton are particularly important in plasticity. Rapid activity-dependent changes in synapse volume or shape are crucial for increasing synaptic strength (Spence and Soderling, 2015). Disruptions of the synaptic cytoskeleton affect the stability and maturation of synapses and subsequently disturb neuronal communication (Goellner and Aberle, 2012). One of the cytoskeleton proteins that were identified in our study is ArpC3. Alterations of ArpC3 activity in excitatory neurons lead to the asymmetric structural plasticity of dendritic spines, followed by the progressive loss of spine synapses (Kim et al., 2013). ArpC3 has been shown to be present in S-PALM and S-NO forms, but the molecular function of these modifications remains unclear (Kang et al., 2008).

Previous studies showed that many enzymes that are involved in cellular energy metabolism, including glycolysis and fatty acid oxidation, are susceptible to modifications by either palmitate or nitric oxide, and some are susceptible to both (Zaręba-Kozioł et al., 2014, Percher et al., 2016). In the present study, we observed chronic stress-induced alterations of PTM crosstalk of the following proteins that are involved in energy metabolism: malate dehydrogenase, citrate synthase, pyruvate dehydrogenase, and short/branched chain specific acyl-CoA dehydrogenase. Palmitate was shown to regulate the activity of such enzymes as malate dehydrogenase and ATP-citrate synthase by affecting the formation of their active ternary complexes (Kostiuk et al., 2008). Stress-induced changes in the interplay between PTMs of proteins that are involved in energy metabolism may decrease the activity of metabolic pathways and thus decrease ATP production.

TCA cycle activation is directly involved in neurotransmission in the brain. Glutamate dehydrogenase S-NO inhibits glutamate oxidation and promotes its conversion to glutamine (Cooper, 2012). Moreover, the absence of S-NO of this enzyme results in a higher ratio of glutamine to glutamate, consistent with the increase in glutamate oxidation (Raju et al., 2015). The association between chronic stress and dysfunction of the mechanisms that regulate glutamate metabolism have been very well described (Sanacora et al., 2012, Popoli et al., 2011). Lower levels of glutamate/glutamine uptake and cycling were found in the brains of chronically stressed mice (Sanacora et al., 2012). Furthermore, higher glutamate levels and lower glutamine/glutamate ratios have been consistently detected in the plasma of depressed patients (Abdallah et al., 2014). The PTM interplay of enzymes that govern the glutamine-glutamate cycle may be involved in these changes.

In each of the functional groups, we can distinguish proteins with kinase and protein phosphatase activity. Among the various PTMs, protein phosphorylation is the most prevalent modification that regulates synaptic protein structures and functions in a wide range of cellular processes (Zahid et al., 2012, Xia et al., 2008, Ubersax and Ferrell, 2007). The dysregulation of protein kinases and phosphatases has been widely reported to be associated with the development of different diseases (Ardito et al., 2017). The reciprocal interplay between S-PALM and phosphorylation was recognized many years ago for the β-adrenergic G protein-coupled receptor (Adachi et al., 2016, Moritz et al., 2015). Recent studies confirmed this molecular mechanism for other neuronal proteins, such as α-amino-3-hydroxy-5-methyl-4-isoxazolepropionic acid and NMDA receptors (Gauthier-Kemper et al., 2014). Our data indicate that S-PALM/S-NO crosstalk may also influence phosphorylation by regulating phosphorylating enzyme machinery.

Identification of the interplay between different types of PTMs of the same protein has increased our understanding of signal processing, but it has not yet been characterized on a larger scale. In the present study, we employed an MS approach, called PANIMoni, that allows the site-specific analysis of two important cysteine modifications: S-PALM and S-NO. We applied this method to identify chronic stress-induced site-specific dynamic changes in PTMs of mouse brain PSD proteins. Our results demonstrated that the S-PALM/S-NO crosstalk of synaptic proteins under physiological conditions modulates almost all aspects of synaptic function. Our data indicate that this interplay is altered by chronic stress. We hypothesize that alterations of S-PALM/S-NO crosstalk of proteins that are involved in synaptic transmission, protein localization, and the regulation of synaptic plasticity might be a major event that leads to the destabilization of synaptic networks in chronic stress-related disorders.

## Methods

### Chemicals

Bradford reagent, sucrose, Ficoll, neocuproine, N-ethylmaleimide, corticosterone, SNAP, L-NAME, 2-bromohexadecanoic acid (2-Br), copper sulfate (CuSO_4_), Tris(2-carboxyethyl)phosphine hydrochloride (TCEP), and sodium ascorbate were purchased from Sigma. Neutravidin-agarose, N-[6-(biotinamido)hexyl]-3’-(2’-pyridyldithio)propionamide (biotin-HPDP), and Oregon Green 488 azide were purchased from Thermo Fisher Scientific. Sequencing-grade modified trypsin was obtained from Promega. Complete protease inhibitor cocktail was obtained from Roche Diagnostics. Enhanced chemiluminescence reagents were purchased from Amersham Biosciences. Alkyne-palmitic acid was obtained from Iris Biotech. The corticosterone enzyme-linked immunosorbent assay (ELISA) kit was purchased from Enzo Life Sciences.

### Chronic restraint stress

Twelve-to 14-week-old wild type male C57/BL6 mice were subjected to restraint stress as previously described (Pawlak et al., 2003). The procedures were performed during the light period of the circadian cycle. All of the animals were individually housed with free access to food and water. The mice were weighed every other day. After the initial 7 days, during which all of the mice were subjected to handling only, control mice were left undisturbed, and stressed animals were subjected to restraint stress for 6 h/day for 21 days in a separate room. The experiments were approved by the Local Bioethical Committee.

### Tail suspension test

One day after the chronic restraint stress procedure, stressed and control mice were subjected to the tail suspension test by hanging them by their tails according to a previously described protocol (Can et al., 2011). The test was recorded, and immobility time was measured for 6 min using a stopwatch.

### Measurement of serum corticosterone

At the end of the experiment (2 days after the last restraint session), stressed and control mice were euthanized by cervical dislocation and decapitated. Blood was collected and centrifuged. Corticosterone levels were measured using an ELISA kit according to the manufacturer’s instructions.

### Synaptoneurosomes isolation

Synaptosomes were prepared from the brains of control and stressed mice as previously describe (Zareba-Koziol et al., 2014). After euthanasia by cervical dislocation, the mice were decapitated. The brains were immediately removed and homogenized using a Dounce homogenizer in 6 ml of buffer A that contained 5 mM HEPES (pH 7.4), 0.32 M sucrose, 0.2 mM ethylenediaminetetraacetic acid (EDTA), 50 mM NEM (a free thiol blocking reagent), and protease inhibitor cocktail. The homogenate was centrifuged at 2,500 × g for 5 min, yielding a pellet and supernatant fractions. The supernatant was then centrifuged at 12,000 × g for 5 min. The pellet was resuspended in buffer A, placed on a discontinuous Ficoll gradient (4%, 6%, and 13%), and centrifuged at 70,000 × g for 45 min. The synaptosomal fraction was collected in buffer A and centrifuged at 20,000 × g for 20 min. The pellet corresponded to the synaptoneurosome fraction.

### Postsynaptic density fraction isolation

Isolated synaptoneurosomes were further diluted with 5 ml of 1% TritonX-100 in 32 mM sucrose and 12 mM Tris-HCL (pH 8.1). The sample was stirred for 15 min in the same open-top tube in a 4°C cold room and then centrifuged at 33,000 × g for 20 min. The pellet was resuspended with 500 µl of buffer solution and layered onto a sucrose gradient that contained 4 ml of 1.5 M sucrose and 4 ml of 1.0 M sucrose. The sample was spun in a swing rotor at 200,000 × g for 2 h. The streak-like cloudy band between 1.0 M sucrose and 1.5 M sucrose that containing PSDs was carefully removed and resuspended in 600 µl of buffer solution. An equal amount of 1% Triton X-100 and 150 mM KCl was added to the sample for resuspension. The sample was then centrifuged at 200,000 × g for 30 min. The resulting pellet contained the PSD fraction.

### Biotin switch method/Acyl Biotin Exchange

The substitution of S-nitrosylated Cys (S-NO-Cys) sites or S-palmitoylated Cys sites with S-biotinylated Cys in PSD protein lysates was based on a previously described biotin switch method and ABE procedure (Jaffrey and Snyder, 2001, Wan et al., 2007). Postsynaptic density protein fractions that were obtained in the previous steps were dissolved in HEN buffer that contained 250 mM HEPES (pH 7.7), 1 mM EDTA, and 0.1 mM neocuproine. To avoid rearrangements of thiol-modifying groups, the protein mixture was treated with blocking buffer solution that contained 250 mM HEPES (pH 7.7), 1 mM EDTA, 0.1 mM neocuproine, 5% sodium dodecyl sulfate (SDS), and 50 mM N-ethylmaleimide at 4°C for 16 h with agitation. To remove excess reagent, the protein extracts were precipitated with 96% of ethanol and resuspended in the same volume of HEN buffer with 2.5% SDS. The obtained protein solutions were then divided into two equal parts. One part was treated with a mixture of 400 µM Biotin-HPDP and 5 mM sodium ascorbate for S-NO and 1 M hydroxylamine for S-PALM. The other part was used as a negative control for the experiments and treated with 400 µM biotin-HPDP without sodium ascorbate or hydroxylamine. All of the samples were incubated in the dark for 1.5 h at room temperature.

### SNO Site Identification (SNOSID)/ Palmitoyl protein identification and site characterization (PalmPISC)

The biotin labeling of S-nitrosylated or S-palmitoylated proteins in lysates was based on previously described procedures (biotin switch method and ABE, respectively). Postsynaptic density protein fractions that contained biotinylated proteins were digested using sequencing-grade modified trypsin (Promega V 5111) for 16 h at 37°C. Digestion was terminated using protease inhibitor cocktail. The tryptic peptide mixture was incubated with 100 μl of neutravidin beads at room temperature for 1 h. The neutravidin beads were washed five times in 1 ml of wash buffer. Neutravidin-bound peptides were eluted with 150 μl of elution buffer (25 mM NH_4_CO_3_ [pH 8.2] and 5 mM TCEP) and concentrated in a SpeedVac. Trifluoroacetic acid was added to the peptide solution to achieve a final concentration of 0.1%. The samples were analyzed by nanoLC-MS and nanoLC-MS/MS

### LC-MS and LC-MS/MS analysis

For each enriched S-PALM peptide-or S-NO peptide-containing sample, separate LC-MS (profile type, peak amplitude data) and LC-MS/MS (peptide identification data) runs were performed. Thermo Orbitrap Elite coupled with Thermo EASY-nLC 1000 was used. S-PALM or S-NO peptides in 0.1% formic acid/water were loaded from a cooled (10°C) autosampler tray to a pre-column (Symmetry C18, 180 µm × 20 mm, 5 µm; Waters) and resolved on a BEH130 column (C18, 75 mm × 250 mm, 1.7 mm; Waters) in a gradient of 5-30% acetonitrile/water for 70 min at a flow rate of 0.3 µl/min. The ultra-performance LC system was directly connected to the ion source of the mass spectrometer. All MS runs were separated by blank runs to reduce the carry-over of peptides from previous samples. All MS runs were performed in triplicate (for both control and stressed mice). The spectrometer resolution was set to 50.000 for MS acquisitions, with an m/z measurement range of 300-2000 Th. Raw LC-MS data were converted to the data format of NMRPipe software (http://spin.niddk.nih.gov/NMRPipe) using an in-house-designed finnigan2Pipe data conversion tool (Bakun et al., 2009, Sikora et al., 2009). MSparky (http://proteom.ibb.waw.pl/mscan/index.html), an in-house-modified version of Sparky NMR software (http://www.cgl.ucsf.edu/home/sparky), was used to convert the LC-MS data into 2D heat maps, in which the peptide’s m/z is one dimension and its LC retention time is the other dimension. MSparky allows interactive validation and inspection of the data. In the LC-MS/MS runs, up to 10 fragmentation events were allowed for each parent ion. Datasets of parent and daughter ions were processed using MascotDistiller 2.5.1 software (MatrixScience). The Mascot search engine (version 2.4.1) was used to survey data against the UniProtKB/Swiss-Prot database (Swissprot 2017_02; 16,905 sequences). The Mascot search parameters were set to the following: taxonomy (Mus musculus), variable modifications (cysteine carbamidomethylation or N-malemideidation, methionine oxidation, peptide tolerance (5 ppm), fragment mass tolerance (0.01 Da). Enzyme specificity was set to trypsin. The lists of the peptide sequences that were identified in all of the LC-MS/MS runs from control and stressed PSD fractions were merged into one selected peptide list (SPL) using in-house-developed MascotScan software (http://proteom.ibb.waw.pl/mscan/index.html). The SPL consists of sequences of peptides with Mascot scores > 20 and unique m/z and LC retention time values. The SPL list of sequences was used to label peaks in all 2D heat maps (profile data) with the same LC, m/z, and z coordinates using in-house-developed TagProfile software. The software allows for the correction of differences in peptide retention times that are caused by changes in the quality of LC columns and slight differences in LC mobile phase content. Acceptance criteria included deviations in m/z values (20 ppm), retention times (10 min), envelope root mean squared error (i.e., a deviation between the expected isotopic envelope of the peak heights and their experimental values, 0.7), and charge state values in accordance with the SPL value. As a result, selected monoisotopic peaks on the 2D heat map were tagged with a peptide sequence. Peptides from the SPL list that were not automatically assigned by TagProfile to appropriate signals on 2D heat maps or those that did not meet all of the acceptance criteria were manually verified using MSparky software. The list of assigned MS sequences was simplified so that a single sequence was assigned for peptides with parent entries that differed only by the charge state (z). The mass spectrometry proteomics data have been deposited to the ProteomeXchange Consortium via the PRIDE partner repository with the dataset identifier PXD010822.

(Reviewer account details: **Username:****; Password:** k9pKB9h4).

### Cortical neuronal culture and treatment

Dissociated cortical cultures were prepared from postnatal day 0 C57/6J mice as previously described (Michaluk et al., 2007). Cells were plated on six-well plates for Western blot or 13-mm-diameter coverslips for S-PALM visualization coated with poly-D-lysine (50 µg/ml) and laminin (2.5 µg/ml) at a concentration of 5 × 10^4^ cells/cm^2^. The cultures were used for the experiments on day 12 in vitro. Corticosterone, 2-Br, and SNAP were dissolved in ethanol. L-NAME was dissolved in water. Cells were pretreated with either corticosterone (1 µM) or 2-Br (30 µM) for 24 h, followed by 30 min exposure to L-NAME (500 µM) or SNAP (100 µM).

### Single-cell in situ imaging of total protein palmitoylation

To visualize the palmitoylated total proteome, neuronal cell cultures were grown on coverslips and incubated overnight with Alkyne-palmitic acid (50 µM) dissolved in dimethylsulfoxide. After washing with phosphate-buffered saline (PBS), the cells were fixed in pre-chilled methanol for 10 min, permeabilized in 0.1% Triton X-100 in PBS, and subjected to click chemistry reaction. Following 1 h incubation at room temperature with click reaction mixture that contained Oregon Green 488 azide (0.1 mM), TCEP (0.1 mM), and CuSO_4_ (0.1 mM), the cells were washed six times with PBS, and the coverslips were mounted in Fluoromount G anti-quenching medium. Images of neurons were acquired using a Zeiss LSM780 laser scanning confocal microscope with a Plan Apochromat 40×/1.4 oil immersion objective.

### Western blot

Cell lysates or total PSD protein fractions after ABE but without neutravidin-based affinity purification were resolved using reducing 10% SDS-polyacrylamide gel electrophoresis. Selectively biotinylated proteins were captured using streptavidin-horseradish peroxidase-conjugated antibodies and visualized by enhanced chemiluminescence (Amersham).

### Functional bioinformatics analysis

Bioinformatics analyses were performed using Panther software, the BINGO plugin in Cytoscape software (release 3.2.0; http://www.cytoscape.org/), and the STRING database. Gene Ontology and REACTOME were performed with 123 proteins that exhibited stress-induced changes in S-PALM/S-NO crosstalk (identified by the PANIMoni method) using the Panther Classification System (version 9.0; www.pantherdb.org). Proteins were classified based on biological process categories in GO annotations. Functional grouping was based on a Fisher Exact test p ≤ 0.05 and at least two counts. As a reference set for term overrepresentation calculations, we utilized proteins that were identified in our MS-measured enrichment analysis and proteins from the Genes to Cognition database. The constructed “PSD reference set” comprised more than 11000 unique mouse proteins. The datasets that were compared are indicated in the respective figures. Enriched GO terms (relative to the our PSD reference set) for selected protein subsets were retrieved using the BINGO Cytoscape plugin and default hypergeometric statistics with Benjamini-Hochberg FDR correction for multiple testing. For the identification of functional annotations, associations, interactions, and networks within our dataset, STRING 9.1 (Search Tool for the Retrieval of Interacting

Genes/Proteins) was used. A threshold confidence level of 0.7 was used to ensure that only highly confident protein interactions were considered for inclusion in the network. Seven types of protein interactions were used for network generation, including neighborhood, gene fusion, co-occurrence, coexpression, experimental, database knowledge, and text mining.

## Acknowledgments

Research was supported by the National Science Centre grant 2015/17/B/NZ3/00557. IF was supported by the National Science Centre grant 2015/19/B/NZ3/01376

## Authors’ contributions

MZK and ABK performed proteomic experiments, analyzed the data; IF performed imaging experiments, analyzed the data and contributed to the manuscript. AK, TW and MB participated in the analysis of data and contributed to the manuscript; MZK and JW conceived the study, designed experiments and wrote the manuscript

All authors read and approved the final manuscript.

